# Evaluation of anti-noroviral compounds on human norovirus replication using 2D monolayers of human intestinal organoids

**DOI:** 10.1101/2025.04.25.650600

**Authors:** Tsuyoshi Hayashi, Yoshiki Fujii, Junki Hirano, Sakura Kobayashi, Kosuke Murakami

**Author notes:** Address correspondence to: Tsuyoshi Hayashi.

## Abstract

We evaluated the anti-noroviral effects of compounds including molnupiravir, on human norovirus (HuNoV) replication using 2D monolayers of human intestinal organoids (HIOs), which have been widely used in HuNoV infection studies. We found that these compounds exhibited lower antiviral activity in 2D HIO monolayers compared with that in apical-out 3D HIOs, a recently developed culture system for HuNoV. This suggests that the anti-noroviral efficacy of these compounds may vary depending on the assay system used.

## Main text

Human norovirus (HuNoV) is a major cause of acute gastroenteritis worldwide, affecting individuals across all age groups (1). HuNoV infection results in symptoms, including nausea, vomiting, and diarrhea. Although typically self-limiting and resolving within a few days to a week in healthy individuals, HuNoV is capable of causing chronic infection in vulnerable populations such as immunocompromised patients. In such cases, it may lead to prolonged viral shedding and gastroenteritis with severe complications (2, 3). Despite its clinical relevance, no antiviral therapies or vaccines are currently available. Efforts to develop anti-HuNoV agents have been limited, partly due to the lack of a reproducible laboratory model for HuNoV replication until recent developments.

In 2016, Ettayebi *et al*. developed a novel HuNoV cultivation system using human intestinal organoids (HIOs) (4). By culturing HIOs as 2D monolayers (See Protocol A in Figure 1), they demonstrated that HuNoVs with diverse genotypes (e.g., GII.3, GII.4, and GII.17) can efficiently replicate in these cells, achieving up to approximately a 3 log_10_-fold increase in viral RNA (4, 5). This cultivation system and infection protocol using 2D HIO monolayers have been widely adopted in norovirus laboratories and are currently used for various HuNoV studies, including those on infection mechanisms, antiviral screening, and virus inactivation (reviewed in references (6, 7)). In antiviral studies, several anti-HuNoV drug candidates have been reported; however, their effects are generally weak to moderate (at *μ*M scale) (Table 1) (7).

**Table 1.**
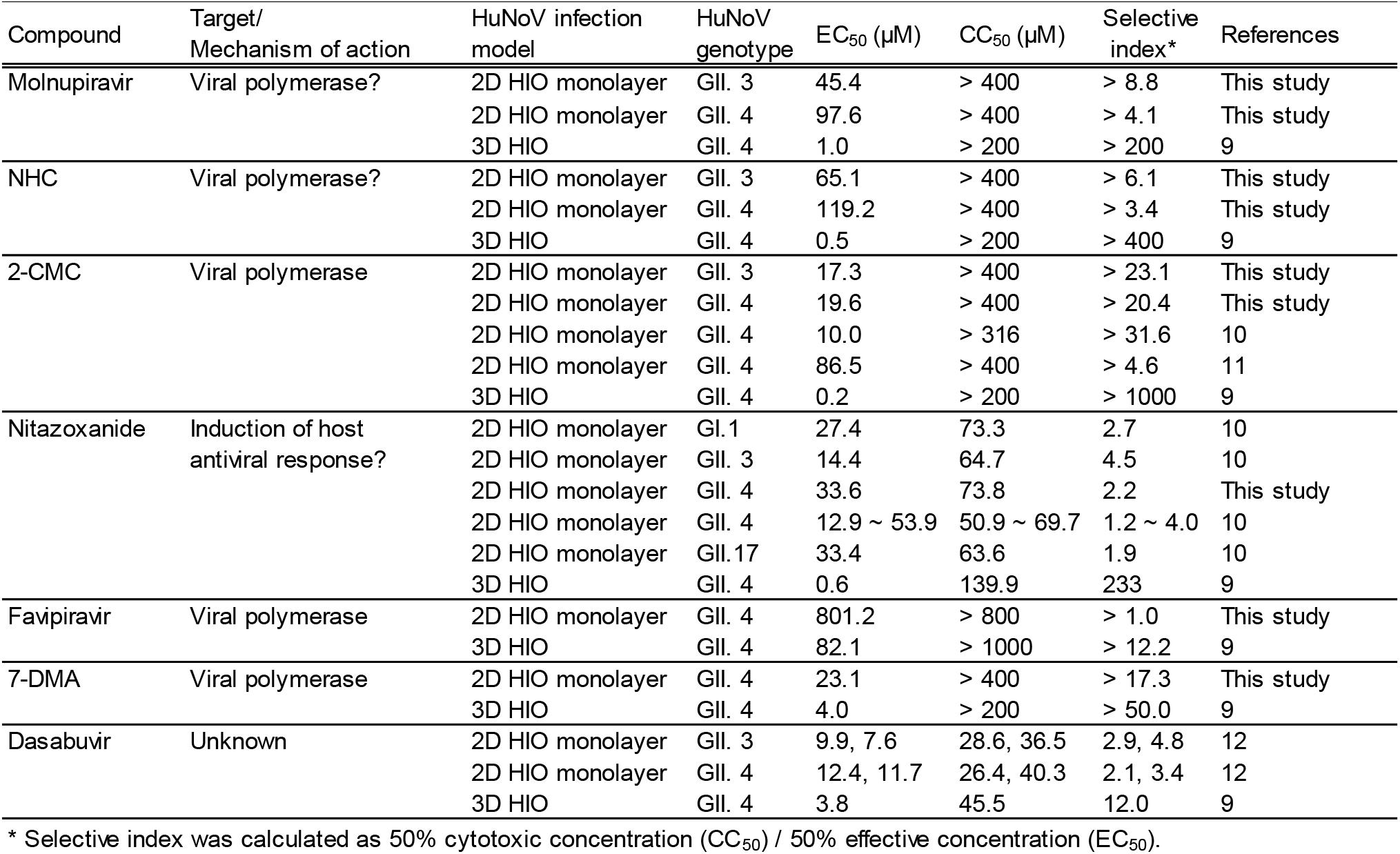
Summary of anti-noroviral compound’s effect assayed by means of 2D HIO monolayers or apical-out 3D HIOs.

**Figure 1.**
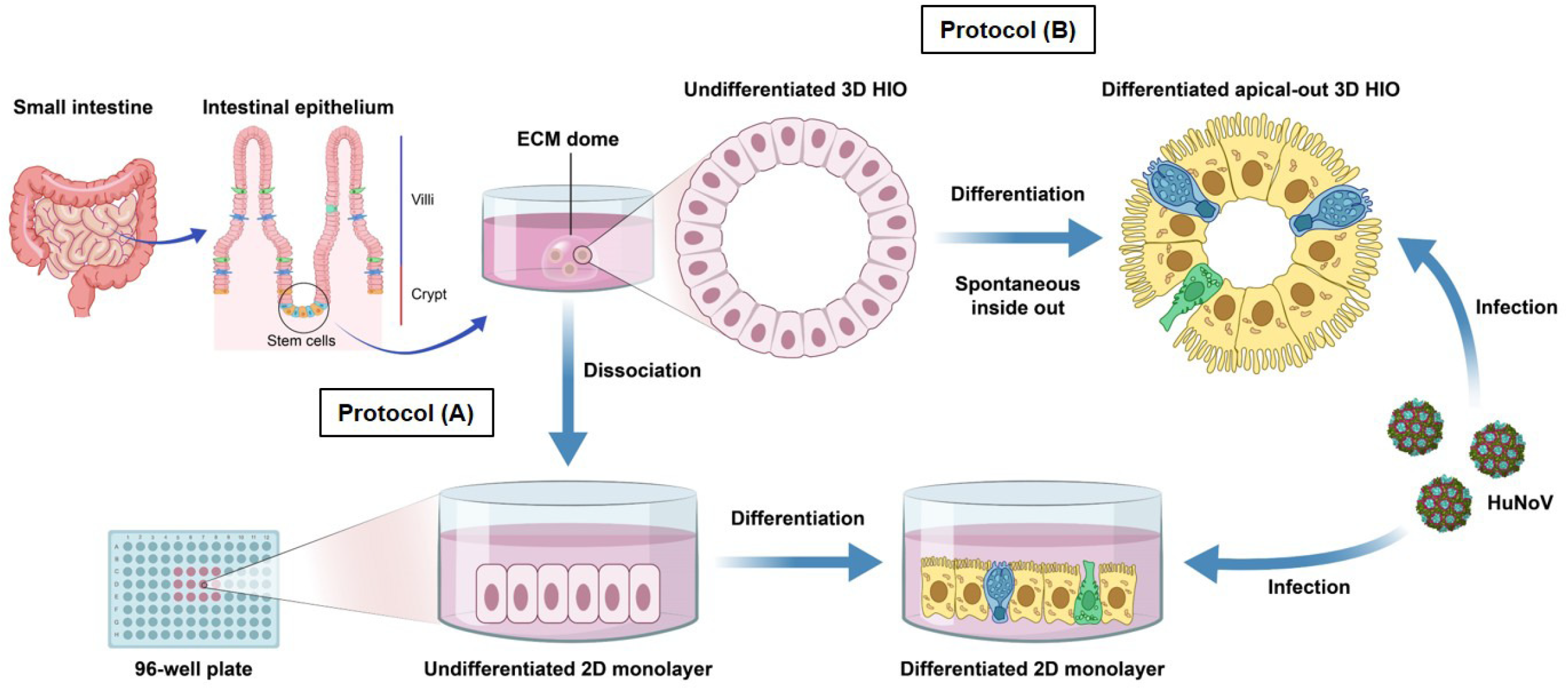
HuNoV infection using 2D HIO monolayers and apical-out 3D HIOs. It is assumed that HuNoV infects host cells via the apical side, which is located inside 3D HIOs. To enhance HuNoV infection efficiency, two protocols have been developed: **Protocol (A):** 3D HIOs are dissociated into single cells, seeded onto a 96-well plate, and cultured as 2D monolayers. Consequently, the apical cell surface is located on the upper side, allowing HuNoV to efficiently bind and infect the cells (5, 14). **Protocol (B);** Mirabelli *et al*. recently reported that during differentiation, 3D HIOs can spontaneously adapt an apical-out orientation, which enables efficient HuNoV infection (8).

More recently, Mirabelli *et al*. developed an alternative HuNoV infection protocol using apical-out 3D HIOs, which reportedly support efficient HuNoV replication (See Protocol B in Figure 1) (8). Using this system, Santos-Ferreira *et al*. demonstrated that molnupiravir, an antiviral known to show inhibitory effects on several RNA viruses (e.g., SARS-CoV-2 and influenza virus) through mutagenesis, strongly inhibits HuNoV replication in 3D HIOs (9). Due to its low 50% effective concentration (EC_50_) (1.0 *μ*M), molnupiravir is considered a promising candidate for anti-HuNoV therapy (9). In this study, the efficacy of other previously reported anti-noroviral compounds, including 2′-C-methylcytidine (2-CMC), nitazoxanide, and dasabuvir, was also tested (9). Interestingly, 2-CMC and nitazoxanide demonstrated over 20-fold greater antiviral activity (lower EC_50_ values) in the 3D HIO system compared with prior results from 2D monolayer assays (Table 1) (10, 11). This discrepancy prompted us to re-evaluate the anti-noroviral effects of these compounds, including molnupiravir, using 2D HIO monolayers.

Differentiated 2D HIO monolayers, derived from 3D HIOs (J2 line), were inoculated with genotype GII.4 or GII.3 HuNoV-positive stool samples in the presence of anti-noroviral compounds in a 100 *μ*L volume. The infection was maintained for 24 h post-inoculation. Subsequently, the cells and 75 *μ*L of the culture medium were harvested for RNA extraction, followed by reverse transcription quantitative PCR (RT-qPCR) analysis to assess viral replication, as previously described (12). The remaining 25 *μ*L of medium was used to evaluate cytotoxicity. The EC_50_ and the half-maximal cytotoxic concentration (CC_50_) for each compound were calculated using GraphPad Prism 9 software.

Consistent with previous findings using apical-out 3D HIOs, molnupiravir and its active metabolite, β-d-N4-hydroxycytidine (NHC), showed dose-dependent inhibition of GII.3 and GII.4 HuNoV replication without compromising cell viability (Figure 2 and Table 1) (9). However, their EC_50_ values were 45.4- to 238.4-fold higher than those reported in 3D HIOs, indicating reduced antiviral efficacy (Table 1). Similar trends were observed for 2-CMC and nitazoxanide (Table 1). In contrast, the difference in EC_50_ for 7-deaza-2′-C-methyladenosine (7-DMA) was relatively small (approximately 6-fold), almost comparable to that of dasabuvir (Table 1). Although its antiviral effect was modest, favipiravir inhibited HuNoV replication in apical-out 3D HIOs (EC_50_ = 82.1 *μ*M; SI > 12.2), whereas its effect was minimal in 2D monolayers (EC_50_ = 801.2 *μ*M; SI > 1.0) (Figure 2, Table 1). These findings suggest that compounds showing weak inhibitory effects in 3D HIOs may have negligible efficacy in 2D monolayer systems.

**Figure 2.**
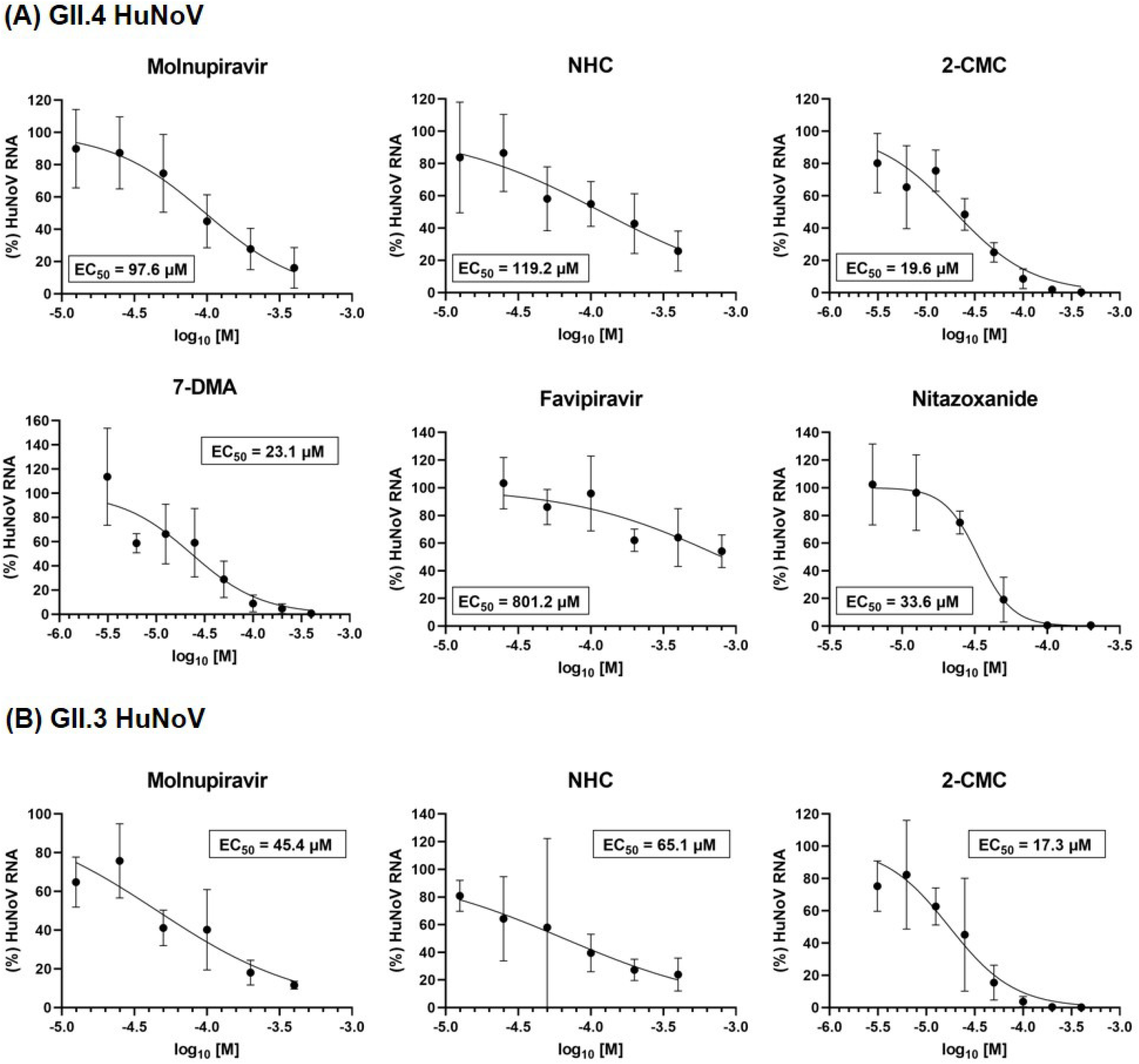
Effect of anti-noroviral compounds on HuNoV replication in 2D HIO monolayers. 2D HIO monolayers were infected with GII.4 or GII.3 HuNoV in the presence of the indicated compounds for 24 h. The cells and supernatants were harvested for RNA extraction, followed by RT-qPCR to quantify HuNoV RNA copies. Experiments were conducted in at least three independent replicates with two or three technical replicates each. Results are normalized to the DMSO control and presented as the mean ± standard deviation (n ≧ 3). EC_50_, 50% effective concentration; CC_50_, 50% cytotoxic concentration.

The reason for the weaker inhibitory effects of the compounds in the 2D monolayers compared with those in 3D HIOs remains unclear. One possibility is that differences in viral replication efficiency may contribute to the variation in antiviral effects. HuNoV replicates more rapidly in 2D monolayers than in 3D HIOs; in 2D monolayers, viral RNA levels increased by 100- to 1,000-fold at 24 h post-infection relative to the input (Figure S1), whereas viral infection in 3D HIOs resulted in approximately 100-fold increases at 48 h (9). Therefore, compounds may have more time to exert inhibitory effects in the context of slower viral replication in 3D HIOs. Additional studies, such as head-to-head comparisons, are needed to elucidate the mechanisms underlying these differences.

A previous study identified molnupiravir as a promising anti-HuNoV candidate (EC_50_ = 1.0 *μ*M in 3D HIOs), while our result indicates that its antiviral efficacy may be overestimated in 3D HIO system. Meanwhile, it should be noted that none of the current HuNoV culture systems, including 2D and 3D HIOs, perfectly reflect acute or persistent HuNoV infections in real-world clinical settings where antiviral treatment would be required. Although significant improvements in the HIO culture system have been achieved through optimized culture media conditions and genetic modifications (5, 13), HuNoV replication efficiency remains lower than that observed during acute infection in humans. In addition, no relevant *in vitro* system mimicking chronic HuNoV infection in humans has been developed, highlighting the need for further optimization and/or development of culture models to more accurately assess anti-noroviral efficacy.

## Supplemental material

**Figure S1. Growth kinetics of HuNoV in 2D HIO monolayers**

2D HIO monolayers were infected with GII.4 or GII.3 HuNoV. At the indicated time points, cells and supernatants were collected to determine viral RNA copy numbers, as shown in Figure 2. Experiments were performed thrice with four technical replicates. Results are presented as the mean ± standard deviation (n =12).

## Author Contributions

Conceptualization, T.H.; methodology, T.H.; software, T.H.; validation, T.H.; formal analysis, T.H.; investigation, T.H., Y.F., S.K., and K.M.; resources, T.H., Y.F., J.H., and K.M.; data curation, T.H.; writing—original draft preparation, T.H.; writing—review and editing, Y.F., J.H., S.K., and K.M.; visualization, T.H. and J.H.; supervision, T.H.; project administration, T.H.; funding acquisition, T.H., Y.F., and K.M. All authors have read and agreed to the published version of the manuscript.

## Acknowledgments

We thank Chizuko Hirano (National Institute of Infectious Diseases, Japan) for technical assistance and Enago (www.enago.jp) for the English language review.

